# Spinal sensorimotor circuits play a prominent role in hindlimb locomotor recovery after staggered thoracic lateral hemisections but cannot restore posture and interlimb coordination during quadrupedal locomotion in adult cats

**DOI:** 10.1101/2023.03.23.533936

**Authors:** Johannie Audet, Sirine Yassine, Charly G Lecomte, Stephen Mari, Soucy Félix, Morency Caroline, Angèle N Merlet, Jonathan Harnie, Claudie Beaulieu, Louis Gendron, Ilya A. Rybak, Boris I. Prilutsky, Alain Frigon

**Author notes:** **Corresponding author** Alain Frigon, Ph.D. Both authors contributed equally (Johannie Audet, Sirine Yassine).

## Abstract

Spinal sensorimotor circuits interact with supraspinal and peripheral inputs to generate quadrupedal locomotion. Ascending and descending spinal pathways ensure coordination between the fore-and hindlimbs. Spinal cord injury disrupts these pathways. To investigate the control of interlimb coordination and hindlimb locomotor recovery, we performed two lateral thoracic hemisections placed on opposite sides of the cord (right T5-T6 and left T10-T11) at an interval of approximately two months in eight adult cats. In three cats, we then made a complete spinal transection caudal to the second hemisection at T12-T13. We collected electromyography and kinematic data during quadrupedal and hindlimb-only locomotion before and after spinal lesions. We show that 1) cats spontaneously recover quadrupedal locomotion following staggered hemisections but require balance assistance after the second one, 2) coordination between the fore-and hindlimbs displays 2:1 patterns and becomes weaker and more variable after both hemisections, 3) left-right asymmetries in hindlimb stance and swing durations appear after the first hemisection and reverse after the second, and 4) support periods reorganize after staggered hemisections to favor support involving both forelimbs and diagonal limbs. Cats expressed hindlimb locomotion the day following spinal transection, indicating that lumbar sensorimotor circuits play a prominent role in hindlimb locomotor recovery after staggered hemisections. These results reflect a series of changes in spinal sensorimotor circuits that allow cats to maintain and recover some level of quadrupedal locomotor functionality with diminished motor commands from the brain and cervical cord, although the control of posture and interlimb coordination remains impaired.

**Significance Statement:** Coordinating the limbs during locomotion depends on pathways in the spinal cord. We used a spinal cord injury model that disrupts communication between the brain and spinal cord by sectioning half of the spinal cord on one side and then about two months later, half the spinal cord on the other side at different levels of the thoracic cord in cats. We show that despite a strong contribution from neural circuits located below the second spinal cord injury in the recovery of hindlimb locomotion, the coordination between the forelimbs and hindlimbs weakens and postural control is impaired. We can use our model to test approaches to restore the control of interlimb coordination and posture during locomotion after spinal cord injury.

## Introduction

Terrestrial locomotion in mammals involves complex dynamic interactions between spinal circuits, supraspinal signals and peripheral sensory inputs [reviewed in (Rossignol et al., 2006; Frigon, 2017; Frigon et al., 2021)]. Musculoskeletal properties also play an important role in stabilizing quadrupedal locomotion and can offset some of loss in neural communication between the brain/cervical cord and the lumbar cord after spinal cord injury (SCI) (Audet et al., 2022). After complete spinal thoracic transection, hindlimb locomotion recovers in various mammals, including mice, rats, cats and dogs (Shurrager and Dykman, 1951; Barbeau and Rossignol, 1987; Bélanger et al., 1996; De Leon et al., 1998, 1999; Cha et al., 2007; Harnie et al., 2019). This recovery involves the locomotor central pattern generator (CPG) that interacts with sensory feedback from the hindlimbs (Brown, 1911; Grillner and Shik, 1973; Grillner and Zangger, 1979; Forssberg et al., 1980; Barbeau and Rossignol, 1987; McCrea and Rybak, 2008; Rossignol and Frigon, 2011; Kiehn, 2016; Grillner and El Manira, 2020; Frigon et al., 2021). Decerebrate cats with a high cervical (C1-C2) transection also express quadrupedal locomotion with pharmacology (Miller and van der Meché, 1976; Miller et al., 1977). However, it is unclear if fore-and hindlimb movements remain coordinated without supraspinal inputs (Frigon, 2017).

Lumbar sensorimotor circuits also play a prominent role in hindlimb locomotor recovery following incomplete SCI (Barrière et al., 2008, 2010). Barriere et al. (2008) performed a dual-lesion paradigm, consisting of a lateral hemisection at T10-T11 followed by complete spinal transection at T12-T13. Instead of taking the minimum 2–3 weeks of treadmill locomotor training usually required, hindlimb locomotion was expressed the day after spinal transection. Thus, after incomplete SCI, plasticity within lumbosacral circuits allowed them to function without motor commands originating from above the spinal transection. The lumbar locomotor CPG likely contributes to hindlimb locomotor recovery after other types of incomplete SCIs.

Another dual spinal lesion paradigm involves performing two lateral hemisections on opposite sides of the cord at different levels (i.e. staggered hemisections) to determine if neural communication remains possible between cervical and lumbosacral levels by activating short propriospinal pathways (Ingebritsen, 1933; Kato et al., 1984, 1985; Stelzner and Cullen, 1991; Courtine et al., 2008; van den Brand et al., 2012; Cowley et al., 2015). (Kato et al., 1984) performed two types of staggered hemisections in adult cats, low thoracic followed by mid-thoracic and high cervical followed by mid-thoracic. In the two types of staggered hemisection paradigms, new fore-hind coordination patterns emerged, with the forelimbs taking more steps than the hindlimbs, or a 2:1 fore-hind coordination, with no consistent phasing between the fore-and hindlimbs during overground locomotion. These results indicate that the spinal locomotor CPGs controlling the forelimbs, located at low cervical/upper thoracic segments (Ballion et al., 2001; Yamaguchi, 2004), operated at a different rhythm and independently from those controlling the hindlimbs, located at upper to mid-lumbar spinal segments (Cazalets et al., 1995; Kiehn and Kjaerulff, 1998; Marcoux and Rossignol, 2000; Kiehn and Butt, 2003; Langlet et al., 2005). However, (Kato et al., 1984) did not separate cycles with 1:1 and 2:1 fore-hind coordination. Studies in intact and single-hemisected cats have shown that step-by-step phasing between the fore-and hindlimbs can remain consistent despite 2:1 coordination during treadmill locomotion (Thibaudier et al. 2013, 2017; Thibaudier and Frigon 2014).

The purpose of the present study was to determine how staggered hemisections affected the control of interlimb coordination and the recovery of hindlimb locomotion. We hypothesize that fore-hind coordination is lost following the second hemisection due to the disruption of direct communication between cervical and lumbar levels. We also hypothesize that spinal sensorimotor circuits play a prominent role in the recovery of hindlimb locomotion following staggered hemisections.

## Materials and Methods

### Ethical approval

The Animal Care Committee of the Université de Sherbrooke approved all procedures in accordance with policies and directives of the Canadian Council on Animal Care (Protocol 442–18). Current data were obtained from eight adult cats (> 1 year of age at the time of experimentation), 4 females and 4 males, weighing between 4.1 kg and 6.5 kg (5.3 ± 1.0). Before and after the experiments, cats were housed and fed in a dedicated room within the animal care facility of the Faculty of Medicine and Health Sciences at the Université de Sherbrooke. Our study followed ARRIVE guidelines for animals studies (Percie du Sert et al., 2020). As part of our effort to reduce the number of animals used in research, all cats participated in other studies to answer different scientific questions, some of which have been published (Lecomte et al., 2022, 2023; Merlet et al., 2022).

### General surgical procedures

Surgical procedures were performed under aseptic conditions with sterilized equipment in an operating room, as described previously (Hurteau et al., 2017; Harnie et al., 2019, 2021; Audet et al., 2022). Before surgery, cats were sedated with an intramuscular injection of butorphanol (0.4 mg/kg), acepromazine (0.1 mg/kg), and glycopyrrolate (0.01 mg/kg). Ketamine/diazepam (0.05 ml/kg) was then injected intramuscularly for induction. Cats were anesthetized with isoflurane (1.5–3%) delivered in O2, first with a mask and then with an endotracheal tube. During surgery, we adjusted isoflurane concentration by monitoring cardiac and respiratory rates, by applying pressure to the paw (to detect limb withdrawal), by assessing the size and reactivity of pupils and by evaluating jaw tone. We shaved the animal’s fur (back, stomach, fore-and hindlimbs) using electric clippers and cleaned the skin with chlorhexidine soap. Cats received a continuous infusion of lactated Ringers solution (3 ml/kg/h) during surgery through a catheter placed in a cephalic vein. A rectal thermometer monitored body temperature, which was maintained within physiological range (37 ± 0.5°C) using a water-filled heating pad placed under the animal and an infrared lamp ∼50 cm over it. At the end of surgery, we injected an antibiotic (Cefovecin, 0.1 ml/kg) subcutaneously and taped a transdermal fentanyl patch (25 mcg/h) to the back of the animal 2–3 cm rostral to the base of the tail to provide prolonged analgesia (4–5-day period before removal). We also injected buprenorphine (0.01 mg/kg), a fast-acting analgesic, subcutaneously at the end of the surgery and a second dose ∼7 h later. Following surgery, we placed the cat in an incubator until they regained consciousness.

### Electrode implantation

We implanted all cats with electrodes to chronically record the electrical activity (EMG, electromyography) of several fore-and hindlimb muscles. We directed pairs of Teflon-insulated multistrain fine wires (AS633; Cooner Wire, Chatsworth, CA, USA) subcutaneously from two head-mounted 34-pin connectors (Omnetics, Minneapolis, MN, USA). Electrodes were sewn into the belly of selected fore-and hindlimb muscles for bipolar recordings, with 1–2 mm of insulation stripped from each wire. We verified electrode placement during surgery by electrically stimulating each muscle through the matching head connector channel. The head connector was secured to the skull using dental acrylic and four to six metallic screws.

### Staggered hemisections and spinal transection

After collecting data in the intact state, a lateral hemisection was made between the fifth and sixth thoracic vertebrae on the right side of the spinal cord. General surgical procedures were the same as described above. The skin was incised between the fifth and sixth thoracic vertebrae and after carefully setting aside muscle and connective tissue, a small laminectomy of the dorsal bone was made. After exposing the spinal cord, we applied xylocaine (lidocaine hydrochloride, 2%) topically and made two to three injections on the right side of the cord. The right side of the spinal cord was then hemisected with surgical scissors between the fifth and sixth thoracic vertebrae. A hemostatic material (Spongostan) was inserted at the lesion site to stop residual bleeding, and muscles and skin were sewn back to close the opening in anatomic layers. In the days following hemisection, cats were carefully monitored for voluntary bodily functions by experienced personnel and bladder and large intestine were manually expressed as needed. The hindlimbs were cleaned as needed to prevent infection. After collecting data following the first hemisection, we performed a second lateral hemisection between the 10^th^ and 11^th^ thoracic vertebrae on the left side of the spinal cord nine to twelve weeks later. Surgical procedures and post-operative care were the same as following the first hemisection. After the second hemisection, we collected data for eight to twelve weeks. In three cats (TO, JA, HO), we performed a complete spinal transection at T12-T13 nine to ten weeks after the second hemisection. We did not perform spinal transections in the other cats because we had to prematurely euthanize them at the start of the covid-19 pandemic. Surgical procedures and post-operative care were the same as following the hemisections.

### Experimental protocol

We collected kinematic and EMG data before (intact state) and at four different time points before and after staggered hemisections during tied-belt (equal left-right speeds) quadrupedal locomotion at 0.4 m/s. The treadmill consisted of two independently controlled running surfaces 120 cm long and 30 cm wide (Bertec, Columbus, OH). A Plexiglas separator (120 cm long, 3 cm high, and 0.5 cm wide) was placed between the left and right belts to prevent the limbs from impeding each other. We present data collected at weeks 1–2 and 7–8 after the first and second hemisections. Cats were not trained to recover quadrupedal locomotion but data collection included several treadmill tasks, such as tied-belt locomotion from 0.4 to 1.0 m/s and split-belt locomotion (left slow/right fast and right slow/left fast), with both the right and left sides stepping on the slow and fast belts (Lecomte et al., 2022). Cats also performed overground locomotion in a straight line and in turns on a custom-built walkway, as well as obstacle negotiations (Lecomte et al., 2023). Some projects also included having cats walk on different surfaces (e.g., foam) to evaluate the influence of somatosensory feedback. We also evoked cutaneous reflexes in some cats by stimulating the superficial radial, superficial peroneal and distal tibial nerves during tied belt and split-belt locomotion at 0.4 m/s and 0.8 m/s. In the intact state and after the first hemisection, nerves were also stimulated with longer trains to induce stumbling corrective reactions in the fore-and hindlimbs during treadmill locomotion at 0.4 m/s and 0.8 m/s (Merlet et al., 2022). Other manuscripts are in preparation. In three cats, we collected data during hindlimb-only locomotion one day, two days, one week, two weeks and three weeks after spinal transection with the forelimbs placed on a stationary platform. At two or three weeks after spinalization, we also collected data during quadrupedal treadmill locomotion at 0.4 m/s in these spinal cats.

In all locomotor trials and at all time points, the goal was to collect ∼15 consecutive cycles using positive reinforcement (food, affection). To avoid fatigue, ∼30 s of rest were given between trials. When required, an experimenter held the tail of the animal to provide mediolateral balance but not to provide weight support. In the double-hemisected and spinal states, some cats, required manual stimulation of the skin of the perineal region to facilitate hindlimb locomotion. For perineal stimulation, the same experimenter manually rubbed/pinched the perineal region with the index finger and thumb. As described, the strength of perineal stimulation is difficult to quantify but we adjusted the pressure applied to the perineal region on a case-by-case basis (light/strong, tonic/rhythmic) to achieve the best hindlimb locomotor pattern possible (Caron et al., 2020; Audet et al., 2022). Perineal stimulation increases spinal excitability and facilitates hindlimb locomotion in spinal mammals through an undefined mechanism(Merlet et al., 2021). However, if the perineal stimulation was too strong, we observed exaggerated flexion of the hindlimbs (hip, knee and ankle) and/or improper left-right alternation, which impaired treadmill locomotion. In other words, too much excitability to spinal locomotor networks was detrimental.

### Data collection and analysis

We collected kinematic and EMG data as described previously (Harnie et al., 2018, 2019, 2021; Lecomte et al., 2021; Audet et al., 2022). Reflective markers were placed on the skin over bony landmarks: the scapula, minor tubercle of the humerus, elbow, wrist, metacarpophalangeal joint and at the tips of the toes for the forelimbs and over the iliac crest, greater trochanter, lateral malleolus, metatarsophalangeal joint and at the tip of the toes for the hindlimbs. Videos of the left and right sides were obtained with two cameras (Basler AcA640–100 g) at 60 frames/s with a spatial resolution of 640 x 480 pixels. A custom-made program (Labview) acquired the images and synchronized acquisition with EMG data. EMG signals were preamplified (10×, custom-made system), bandpass filtered (30–1000 Hz), and amplified (100–5000×) using a 16-channel amplifier (model 3500; A-M Systems). As we implanted more than 16 muscles per cat, we obtained data in each locomotor condition twice, one for each connector, as our data acquisition system is limited to 16 channels. EMG data were digitized (2000 Hz) with a National Instruments card (NI 6032E, Austin, TX, USA), acquired with custom-made acquisition software and stored on computer. In the present study, EMG data are used only for illustrative purposes to show the gait patterns before and after spinal lesions. Measures of EMG and more detailed descriptions will be presented in upcoming papers.

#### Temporal variables

By visual detection, the same experimenter determined, for all four limbs, paw contact as the first frame where the paw made visible contact with the treadmill surface, and liftoff as the most caudal displacement of the toes. We measured cycle duration from successive paw contacts, while stance duration corresponded to the interval of time from foot contact to the most caudal displacement of the toe relative to the hip/shoulder (Halbertsma, 1983). We calculated swing duration as cycle duration minus stance duration. Based on contacts and liftoffs for each limb, we measured individual periods of support (double, triple and quad) and expressed them as a percentage of cycle duration, as described previously (Frigon et al., 2014; Lecomte et al., 2022; Merlet et al., 2022). During a normalized cycle, here defined from successive right hindlimb contacts, we identified nine periods of limb support (Gray and Basmajian, 1968; Wetzel and Stuart, 1976; Frigon et al., 2014; Lecomte et al., 2022). We evaluated temporal interlimb coordination by measuring phase intervals between six pairs of limbs (Thibaudier et al., 2017): 1) left and right forelimbs (forelimb coupling), 2) left and right hindlimbs (hindlimb coupling), 3) left forelimb and left hindlimb (left homolateral coupling), 4) right forelimb and right hindlimb (right homolateral coupling), 5) left forelimb and right hindlimb (right diagonal coupling), and 6) right forelimb and left hindlimb (left diagonal hindlimb). Phase intervals were calculated as the absolute amount of time between contacts of two limbs divided by the cycle duration of the reference limb (English, 1979; English and Lennard, 1982; Orsal et al., 1990; Frigon et al., 2014; Thibaudier and Frigon, 2014; Thibaudier et al., 2017; Audet et al., 2022). The reference limb was always the hindlimb, with the exception of forelimb coupling were it was the right forelimb. For hindlimb coupling, the reference limb was the right hindlimb. Values were then multiplied by 360 and expressed in degrees to illustrate their continuous nature and possible distributions (English and Lennard, 1982; Thibaudier et al., 2017). To determine if single-hemisected and double-hemisected cats displayed greater variations in limb couplings, we calculated the coefficient of variation, a statistical measure of the relative dispersion of data points around the mean, by dividing the standard deviation by the mean, as we described previously (Audet et al., 2022). These values were then multiplied by 100 and expressed as a percentage.

#### Spatial variables

We analyzed spatial variables using DeepLabCut^TM^, an open-source machine learning program with deep neural network (Mathis et al., 2018), as we recently described in the cat (Lecomte et al., 2021). Stride length was measured for the right fore-and right hindlimbs as the distance between contact and liftoff added to the distance traveled by the treadmill during the swing phase, obtained by multiplying swing duration by treadmill speed (Courtine et al., 2005; Goetz et al., 2012; Thibaudier and Frigon, 2014; Dambreville et al., 2015; Lecomte et al., 2021). We measured the relative distance of the paw at contact and liftoff as the horizontal distance between the toe and shoulder or hip markers at stance onset and offset, respectively, for the right fore-and right hindlimbs. As an indicator of limb interference, we measured the horizontal distance between the toe markers of the fore-and hindlimbs on the same side at stance onset and offset of each of the four limbs of the animals.

### Histology and euthanasia

At the end of the experiments, cats were anesthetized with isoflurane before receiving a lethal dose (100 mg/kg) of pentobarbital through the left or right cephalic vein. The extent of the spinal lesion was confirmed by histology, as described previously (Lecomte et al., 2022, 2023). Following euthanasia, a 2 cm length of the spinal cord centered on the lesion sites was dissected and placed in 25 mL of 4% paraformaldehyde solution (PFA in 0.1 m PBS, 4°C). After five days, the spinal cord was cryoprotected in PBS with 30% sucrose for 72 h at 4°C. We then cut the spinal cord in 50 µm coronal sections on gelatinized slides using a cryostat (Leica CM1860, Leica Biosystems Inc, Concord, ON, Canada). Sections were mounted on slides and stained with 1% Cresyl violet. For staining, slides were then dehydrated in successive baths of ethanol 50%, 70% and 100%, 5 minutes each. After a final 5 minutes in a xylene bath, slides were left to dry before being scanned by Nanozoomer (Hamamastu Corporation, Bridgewater Township, NJ, USA). We than performed qualitative and quantitative evaluations of the lesion sites in the transverse plane.

### Statistical analysis

We performed statistical analyses using IBM SPSS Statistics 20.0 software. We first assessed the normality of each variable using the Shapiro Wilk test. As the data were not parametric, we determined the effects of state/time points on dependent variables using the one-factor Friedman test for each state/time points. When a main effect was found, we performed a Wilcoxon signed-rank test with Bonferroni’s correction. The critical level for a statistical significance was set at an α-level of 0.05. Rayleigh’s test was performed to determine whether phase intervals were randomly distributed, as described (Zar, 1974; Kjaerulff and Kiehn, 1996; Thibaudier and Frigon, 2014; Thibaudier et al., 2017; Audet et al., 2022). Briefly, we calculated the r value to measure the dispersion of phase interval values around the mean, with a value of 1 indicating a perfect concentration in one direction, and a value of 0 indicating uniform dispersion. To test the significance of the directional mean, we performed Rayleigh’s z test: z = *n*r^2^, where *n* is the sample size (number of steps). The z value was then compared to a critical z value on Rayleigh’s table to determine if there was a significant concentration around the mean (P value).

## Results

### The recovery of quadrupedal treadmill locomotion after staggered hemisections and extent of spinal lesions

In the present study, all eight cats spontaneously recovered quadrupedal treadmill locomotion at 0.4 m/s one to two weeks following the first lateral hemisection at T5-T6 on the right side. All eight cats also recovered quadrupedal treadmill locomotion at 0.4 m/s one to four weeks following the second lateral hemisection at T10-T11 on the left side. **Figure 1** shows a schematic of the staggered hemisections and the extent of the first and second hemisections for each cat based on histological analysis, which ranged from 40.3% and 66.4% (50.1% ± 9.1) and 33.5% and 53.7% (45.8% ± 6.5) for the first and second lesions, respectively.

**Figure 1.**
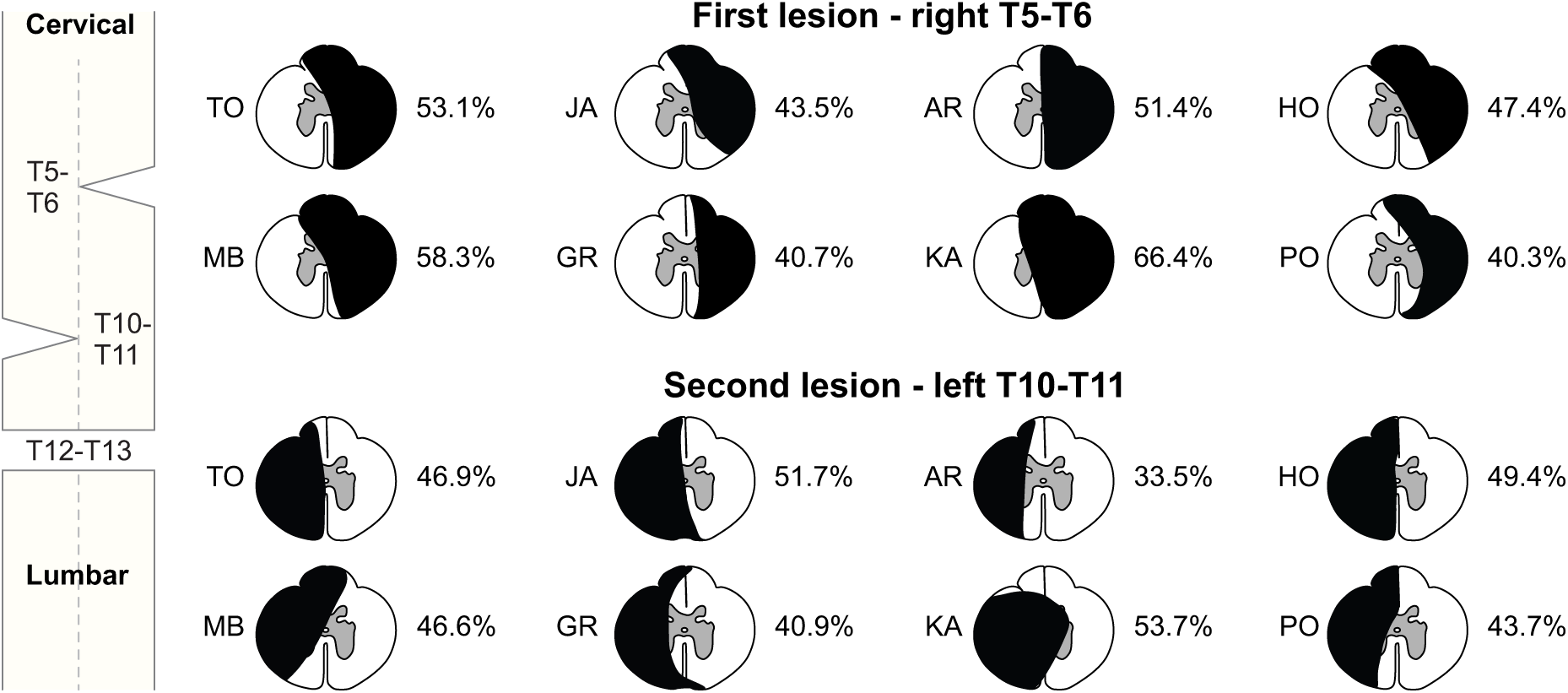
Staggered hemisections paradigm and extent of lesions. Schematic representation of the staggered hemisections and extent of the first and second spinal lesions on the right (T5-T6) and left (T10-T11) sides, respectively, for individual cats. The black area represents the lesioned region.

**Table 1** summarizes three features of locomotor performance after the first and second hemisections. After the first hemisection, only one cat (Cat AR) required balance assistance, where an experimenter held the tail to provide mediolateral balance but not weight support, and only at weeks 1–2. Cats did not require perineal stimulation to perform quadrupedal locomotion after the first hemisection. After the second hemisection, some cats did not recover quadrupedal locomotion until weeks 3 or 4 and all cats required balance assistance at both time points (weeks 1–4 and 7–8). It is important to note that holding the tail was not used for hindquarter weight support, only for balance assistance. After the second hemisection, 5 of 8 and 3 of 8 cats required perineal stimulation at weeks 1–4 and 7–8, respectively. The three cats requiring perineal stimulation at weeks 7–8 also needed it at weeks 1–4.

**Table 1.**
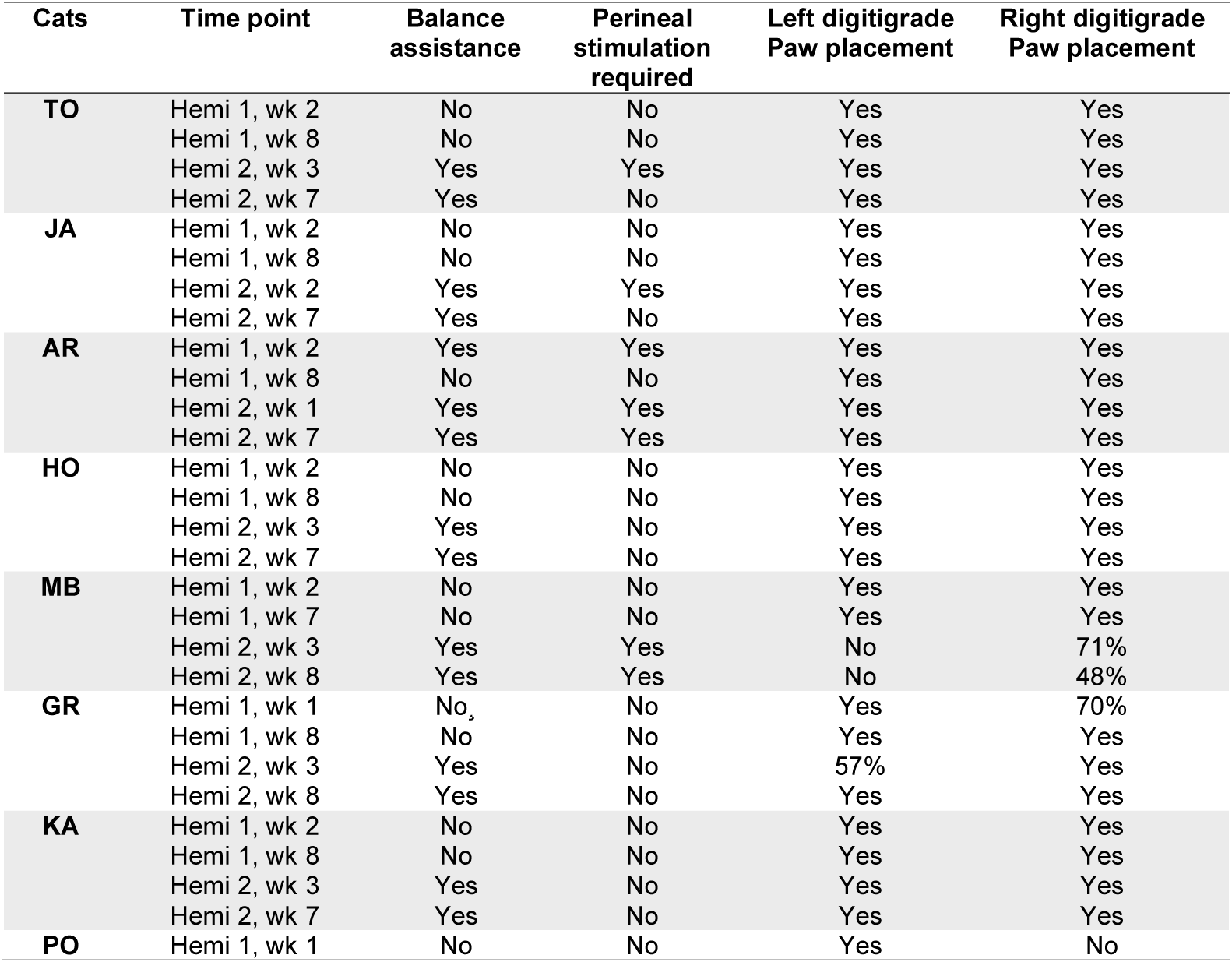
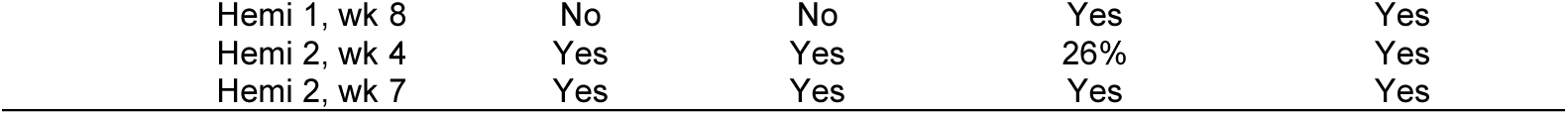
Locomotor performance of individual cats after the first and second hemisections. Locomotor performance of eight cats using four criteria. Percent values indicate the percentage of steps with correct digitigrade placement.

After the first hemisection on the right side of the spinal cord, all eight cats maintained left digitigrade hindpaw placement (contralateral to the lesion). Most cats (6 out of 8) also retained right digitigrade hindpaw placement (ipsilateral to the lesion). However, one cat (Cat PO) showed no digitigrade placement of the right hindpaw at week 1 after the first hemisection, while another cat (Cat GR) performed proper placement 70% of the time. In both cases, the cats placed the right hindpaw on its dorsum. At weeks 7–8 after the first hemisection, all cats performed left and right digitigrade placement. The second hemisection on the left side did not affect digitigrade placement of the right hindpaw in 7 of 8 cats. Only Cat MB showed impaired right hindpaw digitigrade placement with 71% and 48% at weeks 3 and 8, respectively. Surprisingly, most cats (5 out of 8) maintained left digitigrade hindpaw placement at weeks 1–4 after the second hemisection on the left side. Cat MB did not recover left digitigrade placement while cats GR and PO showed impaired left digitigrade placement at weeks 3–4 that recovered at weeks 7–8 after the second hemisection.

### New patterns of forelimb-hindlimb coordination emerge after the first and second hemisections

In the present study, all intact cats performed 1:1 fore-hind coordination in 100% of trials, indicating an equal number of steps at the shoulder and hip girdles, as shown for a single cat in **Figure 2** (top panel). However, at weeks 1–2 and 7–8 after the first hemisection, all cats showed 2:1 fore-hind coordination with varying proportions (48.9% ± 35.4%). When this occurred, cycles with 2:1 and 1:1 fore-hind coordination were intermingled within the same locomotor episode (**Fig. 2**, middle panels) and some cats only showed patterns of 2:1 fore-hind coordination, as shown previously in rats and cats (Górska et al., 1990, 1996, 2013; Bem et al., 1995; Barrière et al., 2010; Alluin et al., 2011; Leszczyńska et al., 2015; Thibaudier et al., 2017). Interestingly, at weeks 1–4 and 7–8 weeks after the second hemisection, some cats displayed a decrease (Cats JA, MB, GR, KA, PO) in the proportion of 2:1 fore-hind coordination while others showed an increase (Cats TO, AR). In cat AR, the proportion of 2:1 coordination increased considerably (**Fig. 2**, bottom panels). **Table 2** summarizes the proportion of 2:1 fore-hind coordination in each cat after both hemisections.

**Figure 2.**
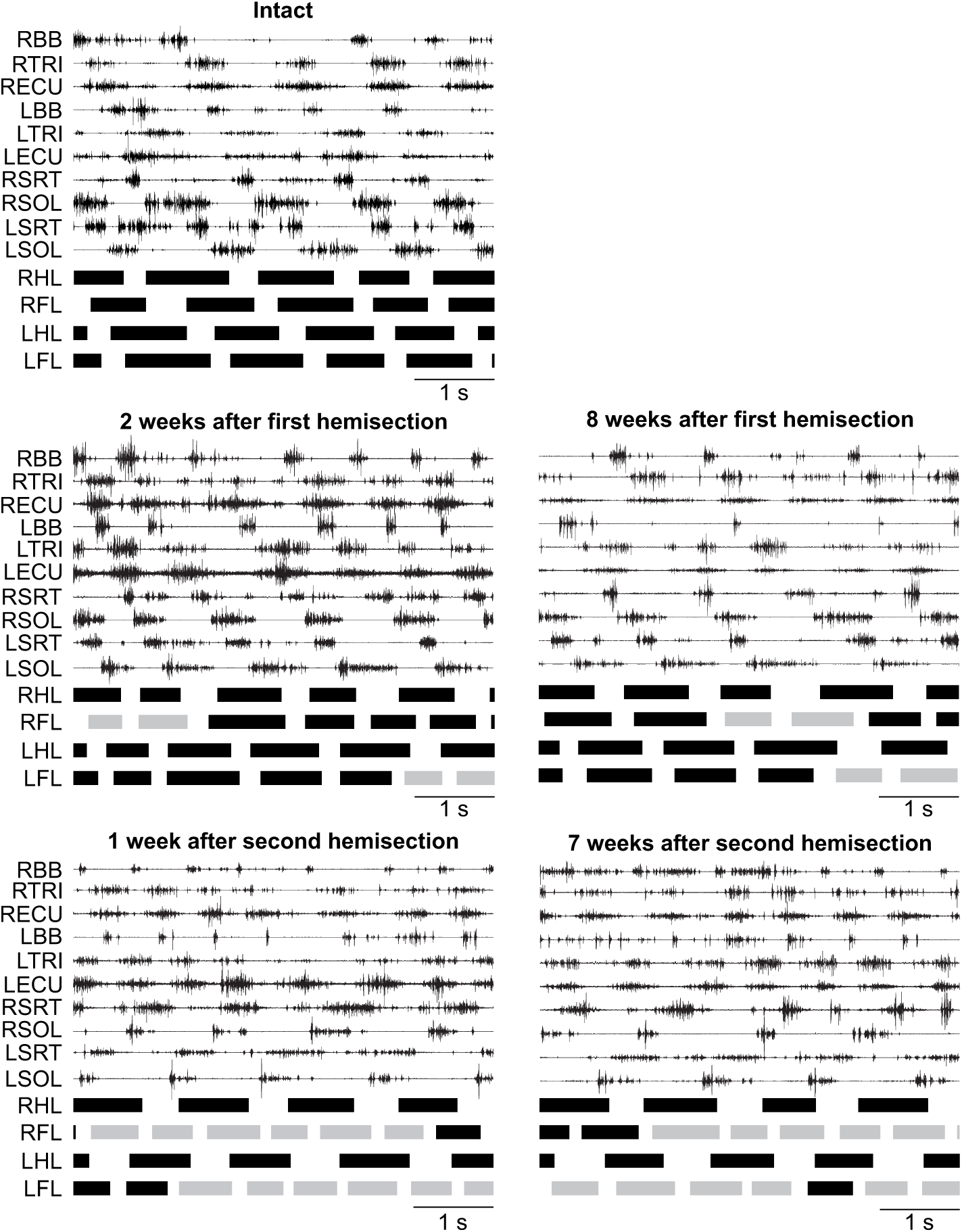
Quadrupedal treadmill locomotion before and after staggered hemisections. Activity from selected fore-(FL) and hindlimb (HL) muscles and stance phases (thick horizontal lines of the left (L) and right (R) limbs in Cat AR at 0.4 m/s. Grey stance phases indicate cycles with 2:1 fore-hind coordination. BB, Biceps brachii; TRI, Triceps brachii; ECU, Extensor carpi ulnaris; SRT, Sartorius; SOL; Soleus.

**Table 2.** Proportion of 2:1 fore-hind coordination after the first and second hemisections. Percent values indicate the percentage of cycles with 2:1 fore-hind coordination while the number in brackets indicate the number of cycles with 2:1 fore-hind coordination dived by the total number of hindlimb cycles recorded.

|  | First hemisection |  | Second hemisection |  |
| --- | --- | --- | --- | --- |
|  | Weeks 1-2 | Weeks 7-8 | Weeks 1-4 | Weeks 7-8 |
| TO | 100% (14/14) | 100% (16/16) | 62% (8/13) | 100% (12/12) |
| JA | 58% (7/12) | 58% (11/19) | 100% (8/8) | 96% (22/23) |
| AR | 50% (4/8) | 100% (21/21) | 75% (18/24) | 96% (23/24) |
| HO | 25% (6/24) | 20% (2/10) | 71% (12/17) | 71% (15/21) |
| MB | 100% (22/22) | 82% (28/34) | 29% (4/14) | 11% (2/19) |
| GR | 10% (1/10) | 19% (7/36) | 52% (11/21) | 50% (6/12) |
| KA | 15% (4/26) | 9% (2/23) | 100% (21/21) | 71% (12/17) |
| PO | 33% (6/18) | 13% (1/8) | 89% (17/19) | 88% (7/8) |

### Interlimb coordination is weaker and more variable after staggered hemisections

To determine how the first and second hemisections affected temporal interlimb coordination, we measured phase intervals between six limb pairs. Values of 0° or 360° indicate a strict in-phase coupling (pacing gait), while a value of 180° indicates a strict out-of-phase coupling. Previous studies in cats have used values between 270° and 90° to denote an in-phase coupling and values between 90° and 270° for out-of-phase coupling (English and Lennard, 1982; Thibaudier and Frigon, 2014; Audet et al., 2022). To assess the step-by-step consistency of forelimb-hindlimb coordination, we performed Rayleigh’s test and calculated the r value, a measure of angular dispersion around the mean for the coupling between the right forelimb and right hindlimb (right homolateral coupling) during tied-belt quadrupedal treadmill locomotion at 0.4 m/s before (intact) and at weeks 1–4 and 7–8 after the first and second hemisections. When the r value is close to 1.0 and significant, it indicates that phase intervals are oriented in a specific direction. We only show this analysis for right homolateral coupling because cats maintained 1:1 coordination between the left and right sides at shoulder (forelimb coupling) and hip (hindlimb coupling) girdles.

In the intact state, we only observed 1:1 fore-hind coordination and right homolateral couplings mainly at 40–80° (**Fig. 3**). At weeks 1–2 and 7–8 after the first hemisection, we found greater dispersal during 1:1 coordination but most right homolateral couplings were from 0–90°. With 2:1 fore-hind coordination, right homolateral couplings were dispersed with the first and second forelimb steps mainly from 0–240° and 120–360°, respectively. After the second hemisection, right homolateral couplings remained dispersed with no clear preference with 1:1 coordination. With 2:1 coordination, right homolateral couplings resembled those found after the first hemisection.

**Figure 3.**
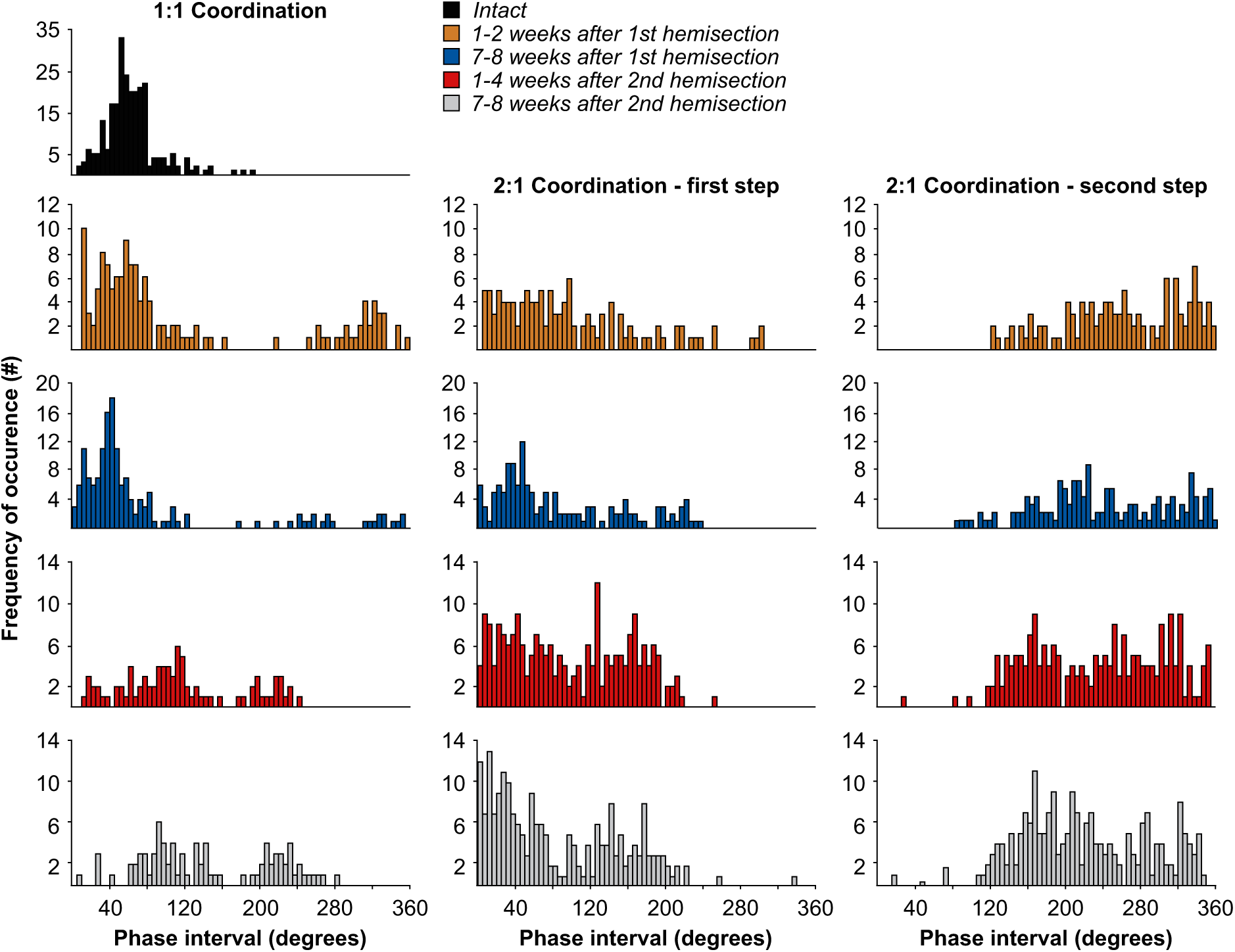
Coordination between right homolateral limbs before and after staggered hemisection. Distribution of right homolateral couplings for the group during 1:1 and 2:1 (first and second forelimb steps) fore-hind coordination. Each bar represents the number of right homolateral couplings found for all eight cats at phase intervals of ten degrees.

**Table 3** shows r values from Rayleigh’s test for phase intervals of right homolateral coupling for individual cats where we separated cycles with 1:1 and 2:1 fore-hind coordination. Note that some cats did not display 1:1 coordination after the first and/or second hemisections. All cats had 1:1 coordination in the intact state, with r values ranging from 0.79 to 1.00 (mean 0.92 ± 0.07). All r values were significant, indicating consistent step-by-step fore-hind coordination. At weeks 1–2 after the first hemisection, six of eight cats had cycles with 1:1 coordination, with r values ranging from 0.45 to 0.94 (mean 0.78 ± 0.21). All r values were significant except for cat JA. At weeks 1–2 after the first hemisection, seven of eight cats had cycles with 2:1 coordination, with r values for the first forelimb step ranging from 0.12 to 0.85 (mean 0.48 ± 0.29). Only cat TO had a significant r value. For the second forelimb step, r values ranged from 0.27 to 0.70 (mean 0.54 ± 0.15) and three r values were significant and four were not. At weeks 7–8 after the first hemisection, six of eight cats had cycles with 1:1 coordination, with r values ranging from 0.13 to 0.85 (mean 0.61 ± 0.26). Four of six cats had significant r values. Six of eight cats had cycles with 2:1 coordination, with r values ranging from 0.30 to 0.84 (mean 0.50 ± 0.23) and 0.12 to 0.80 (mean 0.40 ± 0.25) for the first and second forelimb steps, respectively. Four and three of six cats had significant r vales for the first and second forelimb steps, respectively. At weeks 1–4 after the second hemisection, six of eight cats had cycles with 1:1 coordination, with r values ranging from 0.38 to 0.91 (mean 0.62 ± 0.22). Three of six cats had significant r values. All eight cats had cycles with 2:1 coordination, with r values ranging from 0.39 to 0.83 (mean 0.51 ± 0.14) and 0.12 to 0.86 (mean 0.49 ± 0.21) for the first and second forelimb steps, respectively. Three and four of eight cats had significant r vales for the first and second forelimb steps, respectively. At weeks 7–8 after the second hemisection, only four of eight cats had cycles with 1:1 coordination, with r values ranging from 0.44 to 0.97 (mean 0.72 ± 0.24). Three of four cats had significant r values. All eight cats had cycles with 2:1 coordination, with r values ranging from 0.35 to 0.70 (mean 0.49 ± 0.11) and 0.10 to 0.64 (mean 0.43 ± 0.17) for the first and second forelimb steps, respectively. Three of eight cats had significant r vales for the first and second forelimb steps.

**Table 3.** Circular statistics for forelimb-hindlimb coordination before and after staggered hemisections. The table shows r values from Rayleigh’s test at the different time points for individual cats before and after hemisections for cycles with 1:1 and 2:1 (first and second forelimb steps) coordination. Asterisks indicate a significant r value.

| Cats | Time points | 1:1 coordination | 2:1 – 1<sup>st</sup> step | 2:1 – 2<sup>nd</sup> step |
| --- | --- | --- | --- | --- |
|  |  | r | r | r |
| TO | Intact | 0.79* | - | - |
|  | Hemi 1, wk 2 | - | 0.74* | 0.68* |
|  | Hemi 1, wk 8 | - | 0.75* | 0.44* |
|  | Hemi 2, wk 3 | 0.91* | 0.42 | 0.60* |
|  | Hemi 2, wk 7 | - | 0.70* | 0.64* |
| JA | Intact | 0.86* | - | - |
|  | Hemi 1, wk 2 | 0.45 | 0.64 | 0.27 |
|  | Hemi 1, wk 8 | 0.74* | 0.30 | 0.35 |
|  | Hemi 2, wk 2 | - | 0.83* | 0.86* |
|  | Hemi 2, wk 6 | - | 0.58* | 0.33 |
| AR | Intact | 0.88* | - | - |
|  | Hemi 1, wk 2 | 0.94* | 0.85 | 0.49 |
|  | Hemi 1, wk 8 | - | 0.84* | 0.80* |
|  | Hemi 2, wk 1 | 0.80* | 0.47* | 0.54* |
|  | Hemi 2, wk 7 | - | 0.51* | 0.57* |
| HO | Intact | 0.97* | - | - |
|  | Hemi 1, wk 2 | 0.57* | 0.23 | 0.48 |
|  | Hemi 1, wk 8 | 0.69* | - | - |
|  | Hemi 2, wk 3 | 0.44 | 0.46 | 0.48 |
|  | Hemi 2, wk 7 | 0.87* | 0.42 | 0.42 |
| MB | Intact | 0.96* | - | - |
|  | Hemi 1, wk 2 | - | 0.23 | 0.54* |
|  | Hemi 1, wk 7 | 0.52 | 0.33 | 0.50* |
|  | Hemi 2, wk 3 | 0.71* | 0.39 | 0.49 |
|  | Hemi 2, wk 8 | 0.60* | 0.48 | 0.10 |
| GR | Intact | 1.00* | - | - |
|  | Hemi 1, wk 1 | 0.92* | - | - |
|  | Hemi 1, wk 8 | 0.75* | 0.37 | 0.17 |
|  | Hemi 2, wk 3 | 0.38 | 0.61* | 0.12 |
|  | Hemi 2, wk 8 | 0.44 | 0.41 | 0.39 |
| KA | Intact | 0.97* | - | - |
|  | Hemi 1, wk 2 | 0.93* | 0.12 | 0.64 |
|  | Hemi 1, wk 8 | 0.85* | 0.42 | 0.12 |
|  | Hemi 2, wk 3 | - | 0.46* | 0.38 |
|  | Hemi 2, wk 7 | 0.97* | 0.48 | 0.57* |
| PO | Intact | 0.90* | - | - |
|  | Hemi 1, wk 1 | 0.88* | 0.55 | 0.70* |
|  | Hemi 1, wk 8 | 0.13 | - | - |
|  | Hemi 2, wk 4 | 0.46 | 0.46* | 0.47* |
|  | Hemi 2, wk 7 | - | 0.35 | 0.43 |

Therefore, based on r values and their significance (or lack thereof), as well as the distributions of phase intervals shown in **Figure 3**, fore-hind coordination weakens and becomes more variable after the first hemisection, even when separating cycles with 1:1 and 2:1 coordination. Surprisingly, the second hemisection on the left side had little additional effect on fore-hind coordination compared to what we observed after first hemisection.

To determine how staggered hemisections affected the coordination between limbs of the same girdle, we measured phase intervals for forelimb and hindlimb couplings (**Fig. 4A**). For these analyses, we pooled cycles with 1:1 and 2:1 fore-hind coordination because some cats did not show 1:1 coordination after the first and/or second hemisections. A decrease or an increase in phase interval indicates that the left limb made contact earlier or later, respectively, in the normalized cycle relative to the right limb. For forelimb coupling, we found a significant decrease in the phase interval at weeks 1–4 and 7–8 after the second hemisection compared to the intact state, indicating earlier contact of the left forelimb relative to the right forelimb. For hindlimb coupling, we found a significant increase in the phase interval at weeks 1–4 and 7–8 after the second hemisection compared to weeks 1–2 and 7–8 after the first hemisection, indicating delayed contact of the left hindlimb relative to the right hindlimb.

**Figure 4.**
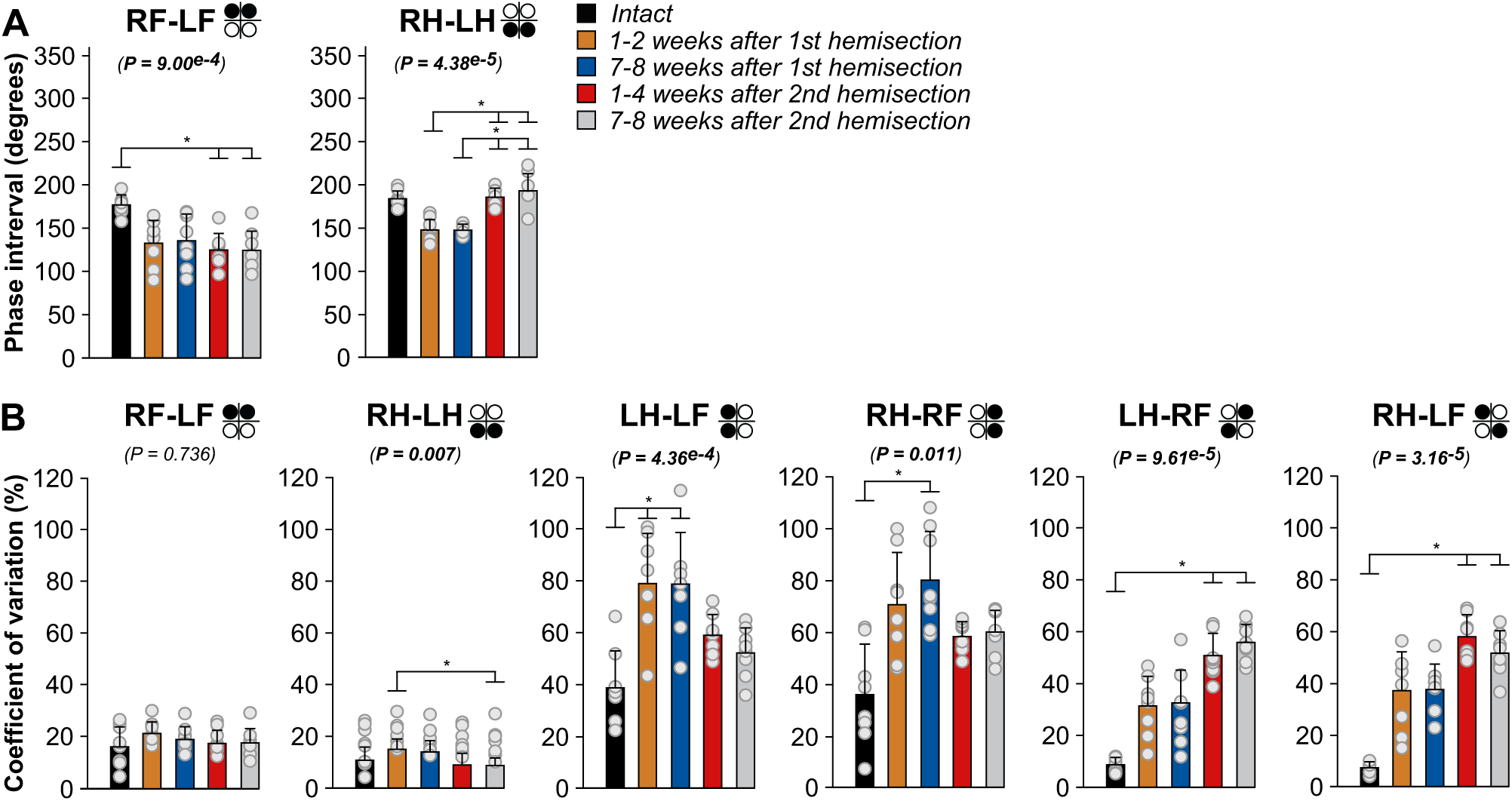
Interlimb phasing and variations during quadrupedal treadmill locomotion before and after staggered hemisections for the group. A) Phase intervals for forelimb and hindlimb couplings. B) Coefficients of variation for six limb pairs. We averaged 8–36 cycles per cat at each time point. The bars represent mean ± SD for the group (n = 8 cats) while grey circles represent individual data points (mean for each cat). The *P* values show the main effect of state (one-factor Friedman test). Asterisks indicate significant differences between time points from the Wilcoxon signed-rank test with Bonferroni’s correction.

To determine if cats displayed greater variations in limb couplings after staggered hemisection, we measured coefficients of variation for all six limb pairs (**Fig. 4B**). We found a significant main effect of state for all limb couplings except for forelimb coupling. Thus, forelimb coupling remains consistent on a step-by-step basis after hemisections. For hindlimb coupling, the coefficient of variation was significantly greater at weeks 7–8 weeks after the second hemisection compared to weeks 1–2 after the first hemisection only. The pattern of change in coefficients of variations for homolateral and diagonal couplings is more revealing. After the first hemisection, we observed a significant increase in the coefficient of variations for left homolateral coupling at both weeks 1–2 and 7–8 compared to the intact state but after the second hemisection, no significant differences with the intact state were found. For right homolateral coupling, the coefficient of variation was significantly greater at weeks 7–8 after the first hemisection compared to the intact state only. Left and right diagonal couplings on the other hand showed greater coefficients of variations after the second hemisection at both weeks 1–4 and 7–8 compared to the intact state. Thus, when considering fore-hind coordination, the second hemisection had no significant additional effect on variability compared to the first hemisection.

### Staggered hemisections generate temporal adjustments in the fore-and hindlimbs and reversals of left-right asymmetries in the hindlimbs

To determine temporal adjustments of the fore-and hindlimbs during quadrupedal treadmill locomotion, we measured cycle and phase durations before and after the two hemisections. For these measurements, we pooled cycles with 1:1 and 2:1 fore-hind coordination because some cats did not show 1:1 coordination after the first and/or second hemisections. For the forelimbs (**Fig. 5A**), we observed a significant reduction in LF and RF cycle and stance durations after the second hemisection at weeks 1–2 and 7–8 compared to the intact state and at weeks 1–4 after the second hemisection compared to weeks 7–8 after the first (**Fig. 5A**). Compared to the intact state, LF and RF swing durations were significantly reduced at weeks 1–4 and 7–8 after the second hemisection, and at weeks 1–4 after the second hemisection compared to weeks 7–8 after the first for LF. Changes in forelimb cycle and phase durations are undoubtedly due to the appearance of 2:1 fore-hind coordination.

**Figure 5.**
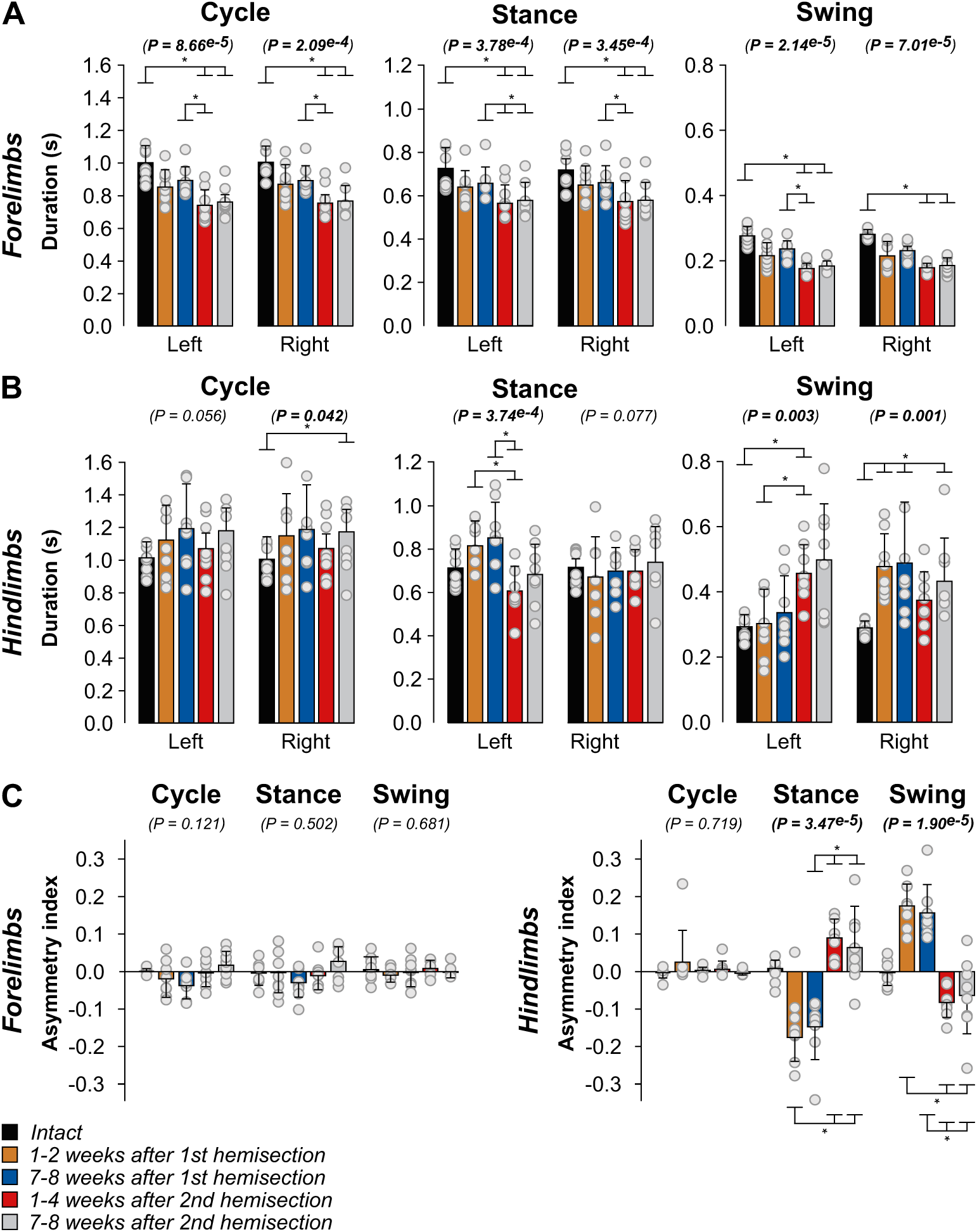
Temporal adjustments during quadrupedal treadmill locomotion before and after staggered hemisections for the group. A and B) Cycle, stance and swing durations for the fore-and hindlimbs, respectively. C) Asymmetry indexes of temporal variables. We averaged 8–36 cycles per cat. The bars represent mean ± SD for the group (n = 8 cats) while grey circles represent individual data points (mean for each cat). The *P* values show the main effect of state (one-factor Friedman test). Asterisks indicate significant differences between time points from the Wilcoxon signed-rank test with Bonferroni’s correction.

For the hindlimbs (**Fig. 5B**), we observed no significant change in LH cycle duration after staggered hemisections, but we found a main effect for RH cycle duration with an increase at weeks 7–8 after the second hemisection compared to the intact state. LH stance duration did not change significantly compared to the intact state after staggered hemisections, but we did observe a significant decrease at weeks 1–4 after the second hemisection compared to weeks 1–2 and 7–8 after the first. RH stance duration did not change significantly after staggered hemisections. LH swing duration was longer at weeks 1–4 after the second hemisection compared to the intact state and weeks 1–2 after the first hemisection. RH swing duration was longer at weeks 1–2 and 7–8 after the first hemisection and at weeks 7–8 after the second hemisection compared to the intact state.

To determine if staggered hemisections produced left-right asymmetries in cycle and phase durations at shoulder and hip girdles, we measured an asymmetry index by subtracting right limb durations from left limb durations (**Fig. 5C**). We found no significant asymmetries in the forelimbs. However, for the hindlimbs, while we observed no asymmetries in cycle duration (cats maintained 1:1 coordination between hindlimbs), stance and swing durations displayed marked asymmetries after the first and second hemisections. The asymmetry index for hindlimb stance duration became negative after the first hemisection, with longer LH stance duration, before switching to positive after the second hemisection, with longer RH stance duration. The asymmetry index for hindlimb swing durations showed an opposite pattern, becoming positive and negative after the first and second hemisections, respectively, indicating that LH swing duration is shorter and longer than RH swing duration after the first and second hemisections, respectively.

### Cats adjust their support periods after staggered hemisections during quadrupedal locomotion

We generally find eight individual support periods during quadrupedal locomotion in a normalized cycle (Frigon et al., 2014; Lecomte et al., 2022). However, a period of double support can become a period of quadrupedal support in some cycles, thus we can find nine different support periods. The proportion of some support periods significantly increased after spinal hemisections, while others decreased (**Fig. 6**). For example, the two periods of triple support involving both hindlimbs (Periods 1 and 5) decreased after the two hemisections compared to the intact state, except at weeks 7–8 after the first hemisection. Periods of diagonal support (Periods 2 and 6) increased after the second hemisection compared to the intact state. Period 2, involving the left forelimb and right hindlimb, increased significantly at weeks 1–4 and 7–8 after the second hemisection compared to the intact state and at weeks 1–4 after the second compared to weeks 1–2 and 7–8 after the first. Period 6 increased at weeks 1–4 and 7–8 after the second hemisection compared to the intact state. The triple support period involving the two forelimbs and the right hindlimb (Period 3) increased after the second hemisection at weeks 1–4 and 7–8 compared to weeks 1–2 and 7–8 after the first. Left homolateral double support (Period 8) did not change significantly after staggered hemisections compared to the intact state. However, it was significantly shorter at weeks 1–4 after the second hemisection compared to both time points after the first hemisection and at weeks 7–8 after the second compared to weeks 7–8 after the first. We observed no significant changes after staggered hemisections for right homolateral support (Period 4), the triple support period involving the left hindlimb and both forelimbs (Period 7) and quadrupedal support (Period 9). Therefore, cats adjust their support periods to maintain dynamic balance during quadrupedal locomotion after staggered hemisections, initially favoring support away from the right hindlimb (side of first hemisection) and then away from the left hindlimb (side of second hemisection).

**Figure 6.**
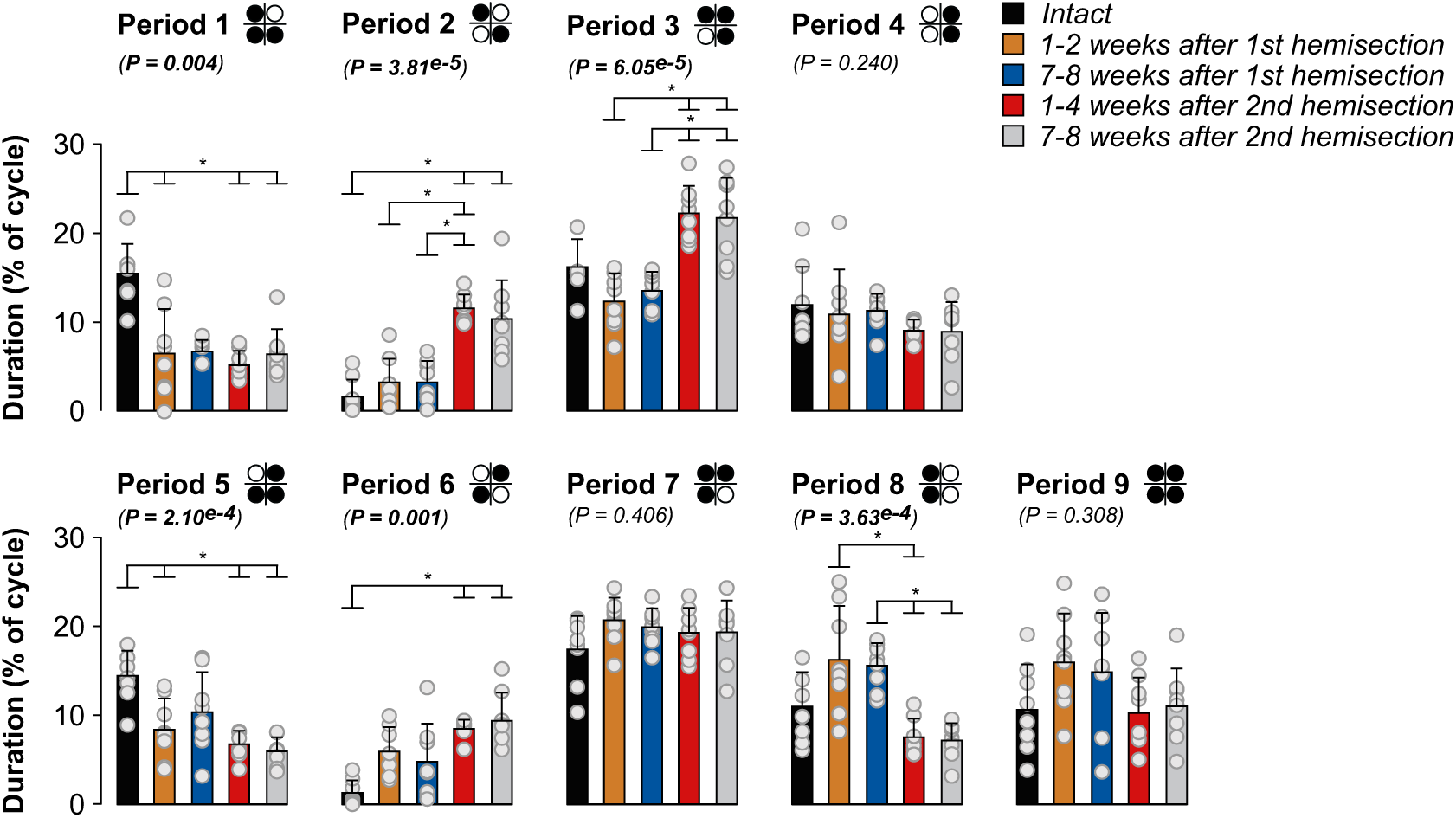
Support periods during quadrupedal treadmill locomotion before and after staggered hemisection for the group. Individual periods of support normalized to right hindlimb cycle duration. We averaged 8–36 cycles per cat. The bars represent mean ± SD for the group (n = 8 cats) while grey circles represent individual data points (mean for each cat). The *P* values show the main effect of state (one-factor Friedman test). Asterisks indicate significant differences between time points from the Wilcoxon signed-rank test with Bonferroni’s correction.

### Staggered hemisections generate spatial adjustments in the fore-and hindlimbs but few left-right spatial asymmetries in the hindlimbs

To determine how staggered hemisections affected spatial parameters, we measured stride length, the horizontal distance traveled by each limb from contact to contact and the horizontal distance of the fore-and hindpaws from the shoulder and hip, respectively, at contact and liftoff. Compared to the intact state, forelimb stride lengths decreased bilaterally but only after the second hemisection, consistent with smaller steps with 2:1 fore-hind coordination (**Fig. 7A**). We observed that the distance of RF relative to the shoulder at liftoff was more rostral at weeks 1–4 after the second hemisection compared to the intact state while LF positioning did not change. Forelimb placement at contact relative to the shoulder did not change significantly.

**Figure 7.**
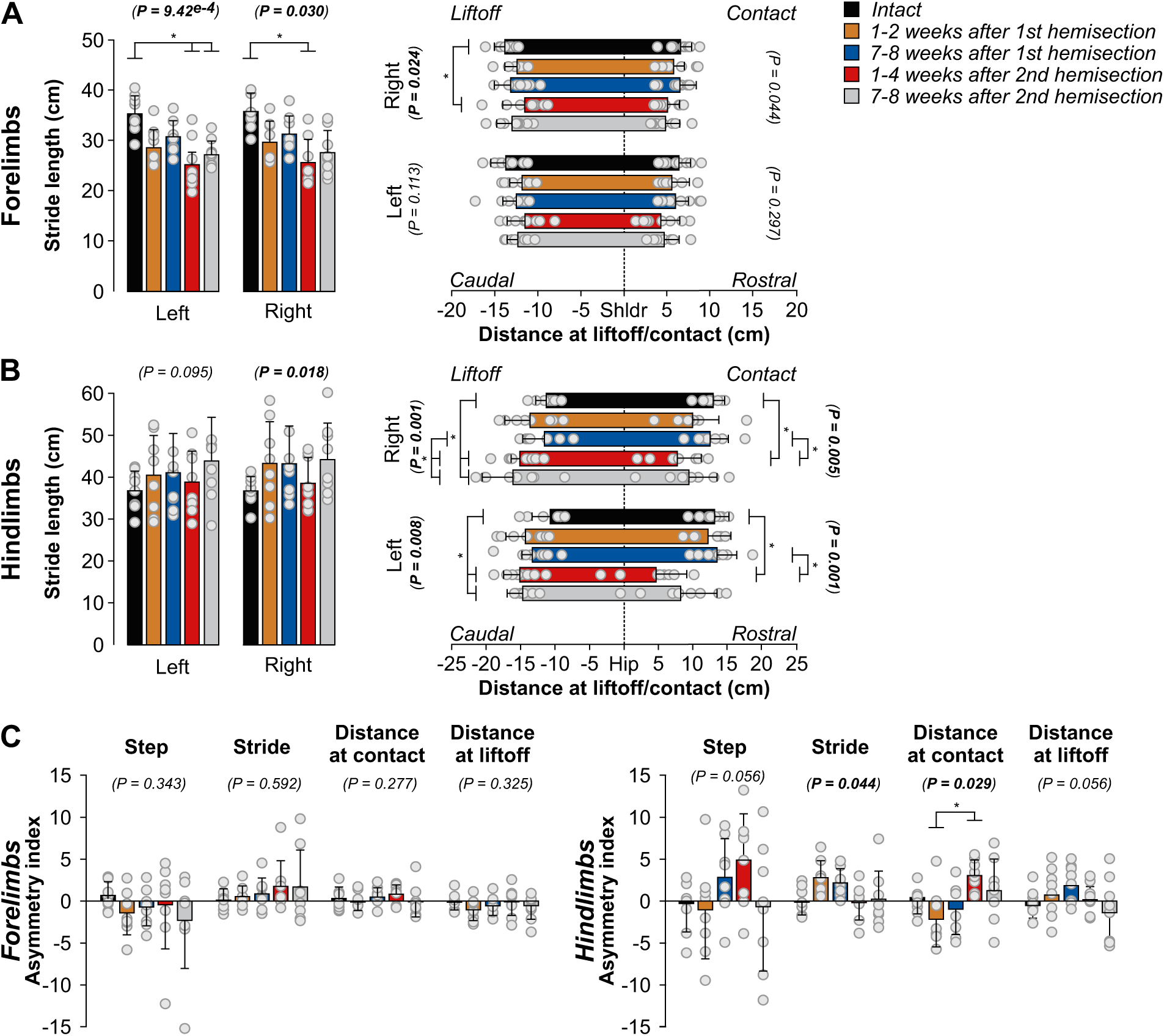
Spatial adjustments during quadrupedal treadmill locomotion before and after staggered hemisections for the group. A and B) Stride length and distances at contact and liftoff for the fore-and hindlimbs, respectively. C) Asymmetry indexes of spatial variables. We averaged 8–36 cycles per cat. The bars represent mean ± SD for the group (n = 8 cats) while grey circles represent individual data points (mean for each cat). The *P* values show the main effect of state (one-factor Friedman test). Asterisks indicate significant differences between time points from the Wilcoxon signed-rank test with Bonferroni’s correction.

Hindlimb stride length did not significantly change after staggered hemisections for LH and although RH showed a significant main effect, we observed no significant difference between time points (**Fig. 7B**). However, we observed several changes in the position of the hindpaw relative to the hip. We found a more caudal horizontal distance between the left hindpaw and the hip at liftoff at both time points after the second hemisection compared to the intact state. Similarly, we found a more caudal horizontal distance between the right hindpaw and the hip at liftoff at both time points after the second hemisection compared to the intact state and at weeks 7–8 after the first hemisection. The right and left hindpaw were closer to the hip at contact at weeks 1–4 after the second hemisection compared to the intact state and weeks 7–8 after the first hemisection.

To determine if staggered hemisections produced asymmetric changes in spatial variables between the left and right sides at shoulder and hip girdles, we measured an asymmetry index by subtracting right limb values from left limb values (**Fig. 7C**). For the forelimbs, we found no significant asymmetries. For the hindlimbs, we found a significant main effect for stride length but pairwise comparisons revealed no differences between time points. For the distance at contact, we only observed a significant difference between weeks 1–2 after the first hemisection and weeks 1–4 after the second hemisection, where left and right placements were more rostral relative to the hip after the first and second hemisections, respectively.

### Forelimb movements adjust to avoid interference after staggered hemisections

To assess limb interference, we measured the horizontal distance between the toe markers of the fore-and hindlimbs at contact and liftoff of the left and right forelimbs (**Fig. 8**), as described previously in spinal cats during quadrupedal locomotion (Audet et al., 2022). The left distance, the distance between LF and LH toe markers, increased at LF contact at weeks 1–4 and 7–8 after the second hemisection compared to the intact state and weeks 7–8 after the first hemisection. At LF liftoff, the left distance increased at weeks 1–4 and 7–8 after the second hemisection compared to the intact state. The right distance, the distance between RF and RH toe markers, increased at weeks 1–2 after the first hemisection and at weeks 1–4 and 7–8 after the second hemisection at both RF contact and liftoff. We propose that increased distances between the fore-and hindlimbs helps avoid interference between the fore-and hindlimbs.

**Figure 8.**
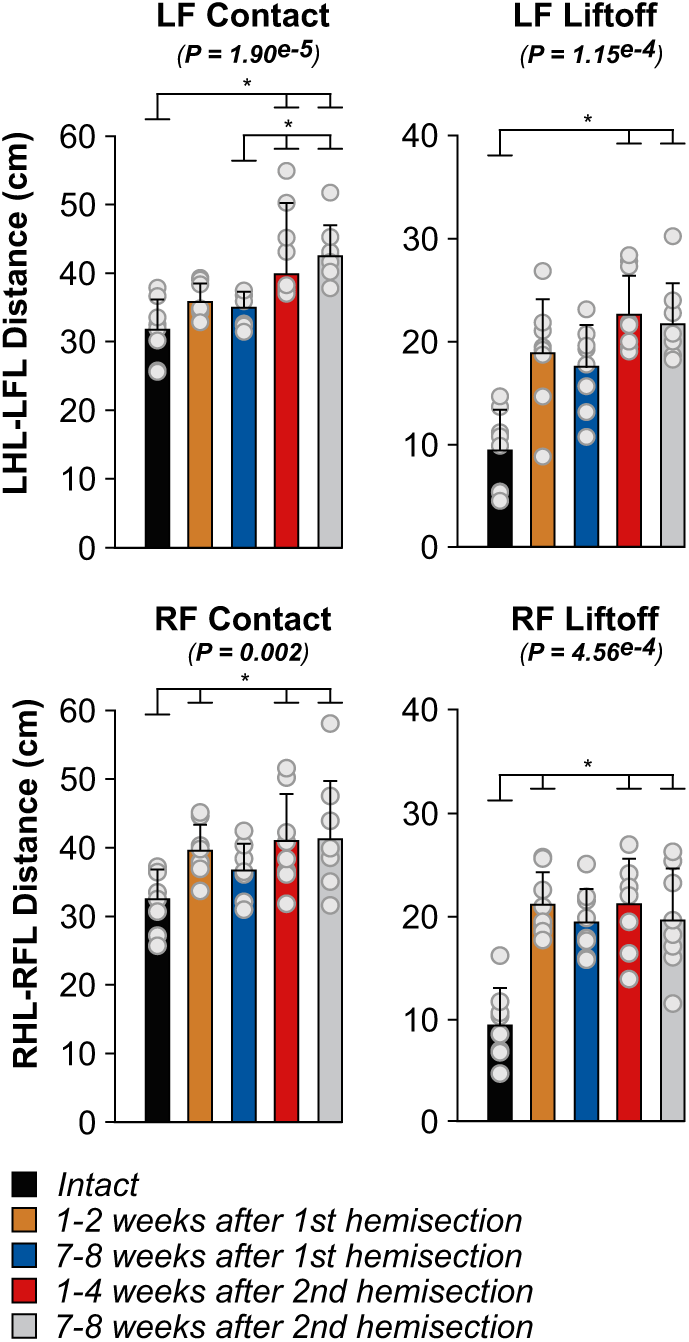
Homolateral limb interference during quadrupedal treadmill locomotion before and after staggered hemisections for the group. Each panel shown horizontal distances between homolateral hindlimbs (HL) and forelimbs (FL) at contact and liftoff of the left and right forelimb. We averaged 8–36 (17.94±7.08) cycles per cat. The bars represent mean ± SD for the group (n = 8 cats) while grey circles represent individual data points (mean for each cat). The *P* values show the main effect of state (one-factor Friedman test). Asterisks indicate significant differences between time points from the Wilcoxon signed-rank test with Bonferroni’s correction.

### The recovery of hindlimb locomotion after staggered hemisections is mediated by a spinal mechanism

As stated in the introduction, a spinal mechanism plays a prominent role in the recovery of hindlimb locomotion following an incomplete SCI (Barrière et al., 2008). To determine if a spinal mechanism also contributes to hindlimb locomotor recovery after staggered hemisections, we performed a spinal transection at T12-T13 nine to ten weeks after the second hemisection in three cats (TO, HO, JA). In all three cats, hindlimb locomotion was expressed the day following transection, a recovery that normally takes a minimum of three weeks (Lovely et al., 1986; Barbeau and Rossignol, 1987; Barrière et al., 2008; Harnie et al., 2019). **Figure 9A** shows a representative example from one cat before transection (i.e. data collected at week 7 after the second hemisection) and at days 1, 2 and 7 after transection without (top panel) and with (bottom panel) perineal stimulation. We can see EMG activity in selected hindlimb muscles during hindlimb-only locomotion. Cat JA stepped one day after the transection without and with perineal stimulation. On the second day, however, the hindlimbs dragged on the treadmill without perineal stimulation but the pattern was robust with perineal stimulation. One week after transection, hindlimb-only locomotion was robust without and with perineal stimulation.

**Figure 9.**
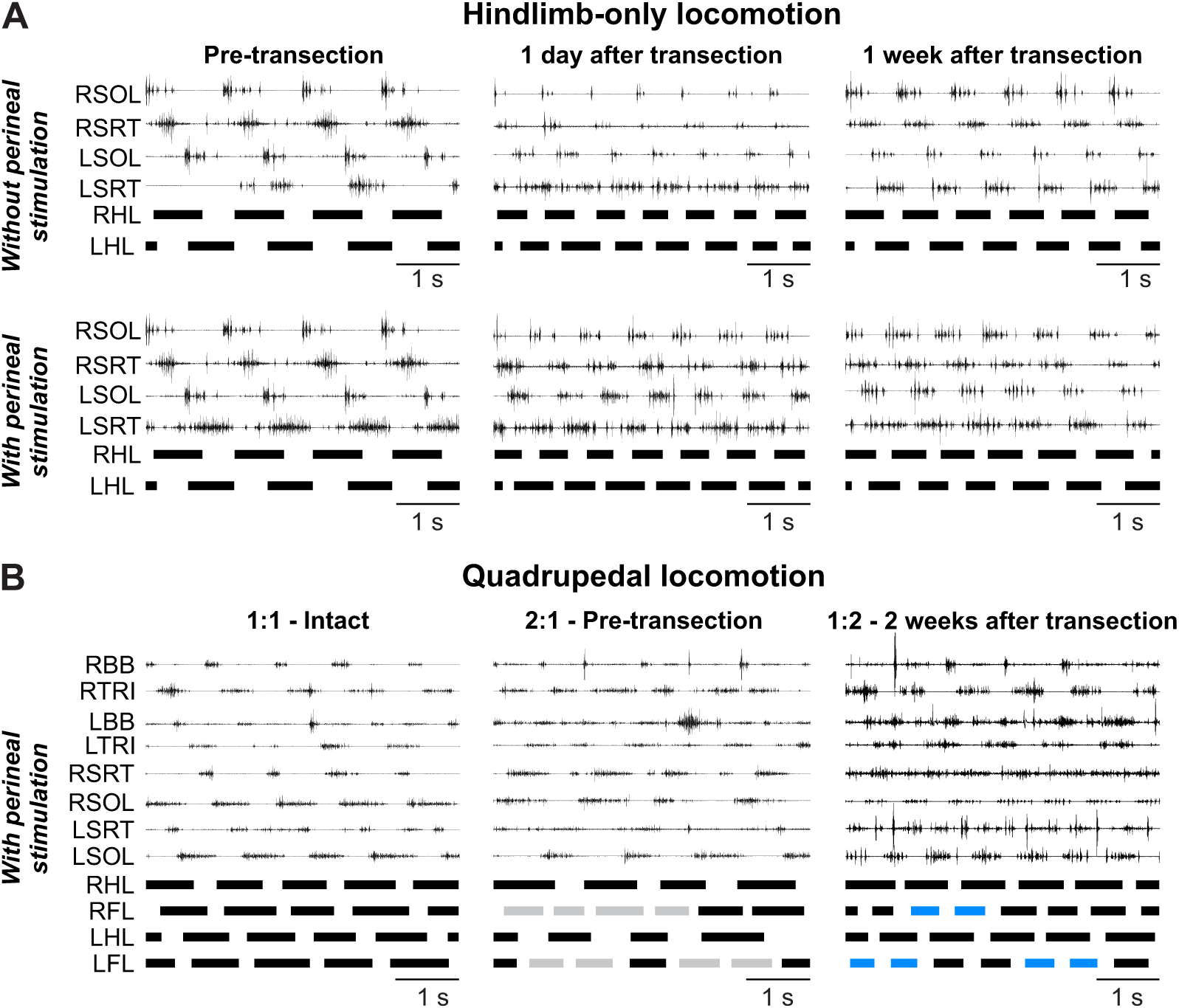
Hindlimb-only and quadrupedal treadmill locomotion before and after complete spinal transection. A) Activity from selected hindlimb muscles and stance phases (thick horizontal lines of the left (LHL) and right (RHL) hindlimbs in Cat JA at 0.4 m/s. B) Activity from selected hindlimb muscles and stance phases (thick horizontal lines of the left (L) and right (R) limbs in Cat HO at 0.4 m/s. Grey and blue stance phases indicate cycles with 2:1 and 1:2 fore-hind coordination, respectively. BB, Biceps brachii; SOL; Soleus; SRT, Sartorius; Triceps brachii.

**Table 4** summarizes three features of locomotor performance before (pre-transection) and at days 1 and 2 as well as weeks 1, 2 after transection without perineal stimulation. Pre-transection, all three cats performed left and right digitigrade paw placement without perineal stimulation but required balance assistance. One day after transection, two cats (TO and HO) did not perform left and right digitigrade paw placement, while Cat JA performed left digitigrade paw placement and right digitigrade paw placement 80% of the time. With the addition of perineal stimulation all three cats performed left and right digitigrade paw placement, but still required balance assistance. Two days after transection, Cats TO performed proper digitigrade placement bilaterally without and with perineal stimulation. In contrast, for Cats JA and HO, perineal stimulation was required to perform proper placement bilaterally. One and two weeks after transection, all three cats performed left and right digitigrade paw placement without and with perineal stimulation, but still required balance assistance. Three weeks after transection, Cat TO only performed proper left digitigrade paw placement 57% of the time without perineal stimulation. In contrast, the two other cats performed proper left and right digitigrade paw placement without and with perineal stimulation.

**Table 4.** Locomotor performance of individual cats before and after spinal transection during hindlimb-only treadmill locomotion without perineal stimulation. Locomotor performance of thre cats using four criteria. Percent values indicate the percentage of steps with correct digitigrade placement.

| Cats | Time point | Balance assistance | Left digitigrade Paw placement | Right digitigrade Paw placement |
| --- | --- | --- | --- | --- |
| TO | Pre spinal | Yes | Yes | Yes |
|  | Spinal, day 1 | Yes | No | No |
|  | Spinal, day 2 | Yes | Yes | Yes |
|  | Spinal, wk 1 | Yes | Yes | Yes |
|  | Spinal, wk 2 | Yes | Yes | Yes |
|  | Spinal, wk 3 | Yes | 57% | Yes |
| JA | Pre spinal | Yes | Yes | Yes |
|  | Spinal, day 1 | Yes | Yes | 80% |
|  | Spinal, day 2 | Yes | No | No |
|  | Spinal, wk 1 | Yes | Yes | Yes |
|  | Spinal, wk 2 | Yes | Yes | Yes |
|  | Spinal, wk 3 | Yes | Yes | Yes |
| HO | Pre spinal | Yes | Yes | Yes |
|  | Spinal, day 1 | Yes | No | No |
|  | Spinal, day 2 | Yes | No | No |
|  | Spinal, wk 1 | Yes | Yes | Yes |
|  | Spinal, wk 2 | Yes | Yes | Yes |
|  | Spinal, wk 3 | Yes | Yes | Yes |

We and others have shown that cats can perform quadrupedal locomotion after a complete thoracic spinal transection (Shurrager and Dykman, 1951; Eidelberg et al., 1980; Howland et al., 1995; Audet et al., 2022). In the present study, all three cats performed quadrupedal treadmill locomotion at 0.4 m/s with perineal stimulation and balance assistance after spinal transection. Instead of 1:1 and 2:1 fore-hind coordination patterns observed at weeks 7–8 after the second hemisection, we observed a 1:2 fore-hind coordination in some cycles, indicating that one hindlimb could take two steps within a single forelimb cycle (**Fig. 9B**), as shown recently in spinal-transected cats (Audet et al., 2022). In all three cats, cycles with 1:2 fore-hind coordination were interspersed with 1:1 coordination. The 1:2 fore-hind coordination represented 21% (Cat TO), 20% (Cat JA) and 48% (Cat HO) of cycles. It is possible that perineal stimulation played a role in the emergence of 1:2 coordination.

## Discussion

We showed that cats spontaneously recovered quadrupedal locomotion following staggered hemisections but required balance assistance after the second. We hypothesized that the second hemisection would more greatly disrupt fore-hind coordination. However, the first hemisection weakened fore-hind coordination and made it more variable, with little additional effect of the second hemisection. Consistent with our hypothesis, hindlimb locomotion was expressed the day after spinal transection in cats that had recovered following the second hemisection. Below we discuss adjustments in the pattern and potential neuroplastic changes that allowed cats to maintain and recover some level of quadrupedal locomotor functionality.

### Recovery of posture and locomotion after staggered hemisections

Lesion extent varied between animals (**Fig. 1**). Generally, smaller lesions associate with faster and more complete locomotor recovery (Barrière et al., 2008; Rossignol et al., 2009). At weeks 1–2 after the first hemisection, only one cat required balance assistance (**Table 1**) while at weeks 7–8, no cat required balance assistance. After the second hemisection, all cats required balance assistance at both time points. Although hindquarter weight support was present in all cats after both hemisections, maintaining posture was challenging after the second. Weight support can be controlled at a spinal level whereas postural control requires supraspinal inputs (Macpherson et al., 1997). Thus, remaining pathways transmitting signals from supraspinal structures and potentially new ones bridging the lesions, such as short propriospinal pathways, are insufficient to restore postural control.

Although all cats recovered quadrupedal locomotion after staggered hemisections, some cats required perineal stimulation after the second hemisection (**Table 1**), which increases spinal neuronal excitability and facilitates hindlimb locomotion in spinal mammals through an undefined mechanism (Eidelberg et al., 1980; Alluin et al., 2015; Harnie et al., 2019; Merlet et al., 2021; Audet et al., 2022). Previous studies proposed that the amount of locomotor training constitutes an important factor in locomotor recovery after partial spinal lesions (Kloos et al., 2005; Rossignol et al., 2009). We recently showed that hindlimb locomotor recovery in spinal cats occurs largely spontaneously without task-specific training (Harnie et al., 2019). Here, although cats did not receive treadmill training after staggered hemisections, they performed various tasks that can be considered training (see Methods). Cats were also freely moved in their cage and in a dedicated room. They could have developed compensatory behavioral strategies through self-training and some cats are naturally more active and athletic than others.

### Interlimb coordination is different, weaker and more variable

We observed 2:1 fore-hind coordination after the first and second hemisections, as shown previously (Eidelberg et al., 1980; Kato et al., 1984; Howland et al., 1995; Jiang and Drew, 1996; Brustein and Rossignol, 1998; Barrière et al., 2010; Alluin et al., 2011; Górska et al., 2013; Thibaudier et al., 2017). Intact cats also perform 2:1 fore-hind coordination on a transverse split-belt treadmill when forelimbs step faster than the hindlimbs (Thibaudier et al., 2013; Thibaudier and Frigon, 2014). This led to the hypothesis that forelimb CPGs have an intrinsically faster rhythmicity than hindlimb CPGs (Thibaudier et al., 2017), which is supported by findings in neonatal rats (Juvin et al., 2005). The 2:1 fore-hind coordination after incomplete SCI could result from reduced inhibition from hindlimb to forelimb CPGs (Górska et al., 2013; Frigon, 2017; Thibaudier et al., 2017), whereby reduced inhibition following thoracic SCI releases the intrinsically faster rhythmicity of forelimb CPGs. Disrupting serotonergic spinal pathways in intact rats also produces 2:1 fore-hind coordination (Sławińska *et al.* 2021). Functionally, 2:1 coordination could represent a strategy to maximize static and dynamic stability (Thibaudier et al., 2017). Performing smaller steps keeps the center of gravity within the support polygon (Cartmill et al., 2002). Another functional reason could be to avoid interference of fore-and hindlimbs (**Fig. 8**). To avoid interference, cats often adopt pacing on a treadmill where homolateral limbs move in phase (Blaszczyk and Loeb, 1993). However, after incomplete SCI, cats might not be able to transition to a pacing gait.

We showed weaker and more variable fore-hind coordination after staggered hemisections (**Figs. 3 and 4**), consistent with previous studies in rats and cats (Kato et al., 1984; Stelzner and Cullen, 1991; Murray et al., 2010; Cowley et al., 2015). The second hemisection did not produce significant additional effects in terms of step-by-step consistency of fore-hind coordination. However, it is important to note that cats required balance assistance after the second hemisection and providing this aid undoubtedly facilitated fore-hind coordination. Impaired coordination between the fore-and hindlimbs could be due to lesioned propriospinal pathways between cervical and lumbar levels and direct supraspinal pathways to the lumbar cord (Sherrington and Laslett, 1903; English, 1980; Kato et al., 1984; Bareyre et al., 2004; Courtine et al., 2008). The loss of interlimb reflex pathways also could have contributed to impaired fore-hind coordination (Hurteau et al., 2018). (Frigon, 2017) argued that fore-hind coordination requires supraspinal commands.

Support periods reorganized after staggered hemisection (**Fig. 6**). Periods of triple support involving the two hindlimbs decreased after the first hemisection and remained decreased after the second. Periods of triple support involving the right hindlimb and both forelimbs, and both diagonal support periods increased after the second hemisection. The cat is most unstable in diagonal support, but these phases help propel the body forward, increasing quadrupedal locomotion efficiency (Farrell et al., 2014). When both forelimbs contact the ground, they provide greater stability. Thus, increased diagonal support and triple support involving the forelimbs could be a strategy to facilitate forward movement while maintaining stability after staggered hemisections.

### Spinal sensorimotor circuits play a prominent role in hindlimb locomotor recovery

Many mammals recover hindlimb locomotion after complete spinal transection because the spinal locomotor CPG can still interact with sensory feedback from the hindlimbs (Shurrager and Dykman, 1951; Lovely et al., 1986, 1990; Barbeau and Rossignol, 1987; Bélanger et al., 1996; De Leon et al., 1998, 1999; Leblond et al., 2003; Cha et al., 2007; Harnie et al., 2019). (Barrière et al., 2008) also showed that the spinal locomotor CPG makes an important contribution to hindlimb locomotor recovery following incomplete SCI. Here, we extend these results by showing that hindlimb locomotion was expressed the day following a spinal transection made 9–10 weeks after the second hemisection (**Fig. 9**). This indicates that the spinal network controlling the hindlimbs had already undergone plastic changes after staggered hemisections, making it more independent from descending signals originating above the lesions. Changes in the spinal cord can include intrinsic changes in neuronal excitability (Murray et al., 2010) and/or in sensorimotor interactions from peripheral afferents (Frigon et al., 2009; Gossard et al., 2015). (Kato et al., 1984) observed that hindlimb movements were initiated following forward movement induced by the forelimbs after staggered hemisections, much like a pantomime horse. Signals from muscle and/or cutaneous afferents likely play a major role in initiating hindlimb movements after staggered hemisections. This is not to say that descending signals cannot still influence and control the lumbar CPG through new short relay propriospinal pathways (Cowley et al., 2015).

### Locomotor recovery involves a series of neuroplastic changes

As mentioned above, we observed several changes in the locomotor pattern. **Figure 10** schematically illustrates potential changes in spinal sensorimotor circuits after staggered hemisections involved in locomotor recovery based on left-right asymmetries in cycle and phase durations (**Fig. 5**) and the immediate expression of hindlimb locomotion after spinal transection. After the first hemisection, ipsilesional lumbar neurons have weaker activity and longer stance phases and increased weight support of the left hindlimb increases load feedback from extensors and cutaneous afferents. The left spinal network increases its influence on the right spinal network. Anatomical and functional asymmetric changes take place within the spinal cord (Murray and Goldberger, 1974; Hultborn and Malmsten, 1983; Helgren and Goldberger, 1993; Frigon et al., 2009). New descending and ascending pathways also form to facilitate descending commands from and to the brain (Fouad et al., 2000; Raineteau et al., 2002; Ballermann and Fouad, 2006; Courtine et al., 2008; Ghosh et al., 2010; Rosenzweig et al., 2010). However, these are insufficient to restore fore-hind coordination. After the second hemisection, neurons of the right spinal network have recovered their activity and stance and weight support is longer/increased for the right hindlimb. The right spinal network increases its influence on the left one. Direct ascending and descending pathways are disrupted but new pathways can form through short propriospinal relays (Zaporozhets et al., 2006; Cowley et al., 2008). However, these are insufficient to restore postural control. Over time after the second hemisection, spinal neuronal activity controlling the left hindlimb recovers. After spinal transection, both left and right spinal networks function without descending inputs and hindlimb locomotion is expressed, possibly via strengthened sensorimotor interactions bilaterally.

**Figure 10.**
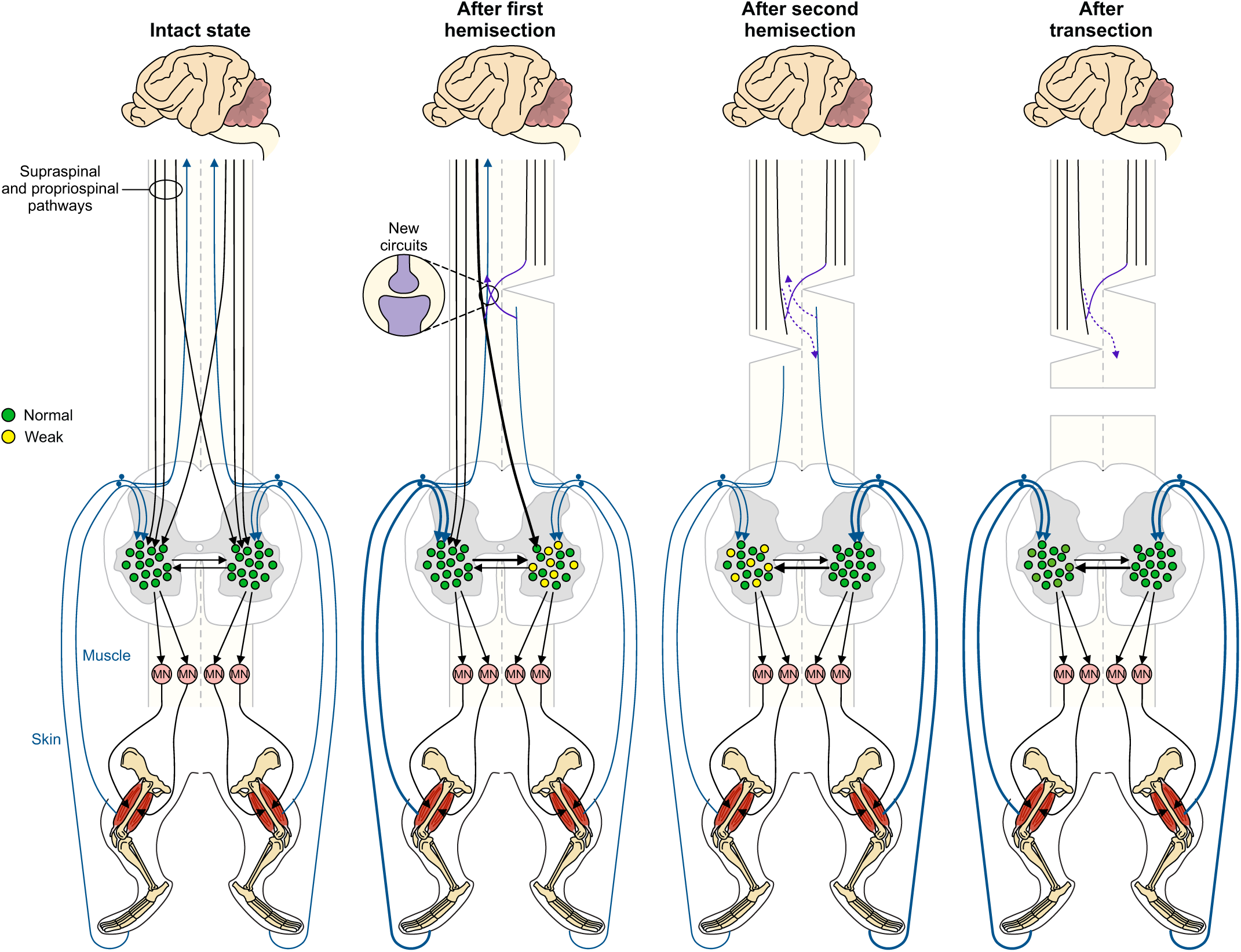
Potential changes in spinal sensorimotor circuits after staggered hemisections. In the intact state, descending supraspinal and propriospinal pathways reach lumbar spinal interneurons that control spinal motoneurons. Pathways transmitting signals from proprioceptive and cutaneous afferents ascend to the brain and project locally to spinal interneurons. After the first hemisection performed on the right side, ipsilesional lumbar neurons have weaker activity and increased weight support of the contralesional hindlimb increases load feedback from extensors and cutaneous afferents. Thicker lines represent increase influence. The left spinal network increases its influence on the right spinal network. New descending and ascending pathways also form to facilitate communication between the brain and spinal cord. After the second hemisection performed on the left side, neurons of the right spinal network have recovered their activity. Direct ascending and descending pathways are disrupted but new pathways can form through short propriospinal relays. After spinal transection, both the left and right spinal network function without descending inputs and hindlimb locomotion is expressed, possibly via strengthened sensorimotor interactions bilaterally.

## Concluding remarks

Staggered hemisections constitute an interesting SCI paradigm to investigate the recovery of posture, interlimb coordination and locomotion. We are currently investigating interlimb reflexes after staggered hemisections and their contribution to postural and locomotor recovery. Future studies need to determine what ascending and descending signals can be transmitted through such lesions, and importantly, if they make meaningful contributions to locomotion and how we can facilitate them using therapeutic approaches.

## Acknowledgments

We thank Philippe Drapeau for providing data acquisition and analysis software, developed in the Rossignol and Drew laboratories at the Université de Montréal. This work was supported by grants from the Natural Sciences and Engineering Research Council of Canada (NSERC RGPIN-2016–03790) to A.F., and the National Institutes of Health: R01 NS110550 to A.F, IAR and BIP. A.F. is a Fonds de Recherche-Santé Quebec (FRQS) Senior Research Scholar. J.H. and A.N.M. were supported by FRQS doctoral and postdoctoral scholarships, respectively. J.A. was supported by master’s scholarships from NSERC and FRQS. C.B. was supported by a master’s scholarship from the Canadian Institutes of Health Research.

